# Altered methionine metabolism impacts phenylpropanoid production and plant development in *Arabidopsis thaliana*

**DOI:** 10.1101/2023.05.29.542770

**Authors:** Doosan Shin, Veronica C. Perez, Gabriella K. Dickinson, Haohao Zhao, Ru Dai, Breanna Tomiczek, Keun Ho Cho, Ning Zhu, Jin Koh, Alexander Grenning, Jeongim Kim

**Affiliations:** Horticultural Sciences Department, University of Florida, Gainesville, FL, 32611; Plant Molecular and Cellular Biology Graduate Program, University of Florida, Gainesville, FL, USA; Genetic Institute, University of Florida, Gainesville, FL, USA; Department of Chemistry, University of Florida, Gainesville, FL, 32611; Interdisciplinary Center for Biotechnology Research, University of Florida, Gainesville, FL, 32611

**Author notes:** Corresponding Author: Jeongim Kim. These authors contributed equally to this work (Doosan Shin, Veronica C. Perez).

**Keywords:** Aldoximes, Phenylpropanoids, Aliphatic glucosinolates, Aliphatic aldoximes, Methionine, Growth and Development, *Arabidopsis thaliana*

## Abstract

Phenylpropanoids are specialized metabolites derived from phenylalanine. Glucosinolates are defense compounds derived mainly from methionine and tryptophan in Arabidopsis. It was previously shown that the phenylpropanoid pathway and glucosinolate production are metabolically linked. The accumulation of indole-3-acetaldoxime (IAOx), the precursor of tryptophan-derived glucosinolates, represses phenylpropanoid biosynthesis through accelerated degradation of phenylalanine-ammonia lyase (PAL). As PAL functions at the entry point of the phenylpropanoid pathway which produces indispensable specialized metabolites such as lignin, aldoxime-mediated phenylpropanoid repression is detrimental to plant survival. Although methionine-derived glucosinolates in Arabidopsis are abundant, any impact of aliphatic aldoximes (AAOx) derived from aliphatic amino acids such as methionine on phenylpropanoid production remains unclear.

Here, we investigate the impact of AAOx accumulation on phenylpropanoid production using Arabidopsis aldoxime mutants, *ref2* and *ref5*. REF2 and REF5 metabolize aldoximes to respective nitrile oxides redundantly, but with different substrate specificities. *ref2* and *ref5* mutants have decreased phenylpropanoid contents due to the accumulation of aldoximes. As REF2 and REF5 have high substrate specificity toward AAOx and IAOx respectively, it was assumed that *ref2* accumulates AAOx, not IAOx. Our study indicates that *ref2* accumulates both AAOx and IAOx. Removing IAOx partially restored phenylpropanoid production in *ref2*, but not to the wild-type level. However, when AAOx biosynthesis was silenced, phenylpropanoid production and PAL activity in *ref2* were completely restored, suggesting an inhibitory effect of AAOx on phenylpropanoid production. Further feeding studies revealed that the abnormal growth phenotype commonly observed in Arabidopsis mutants lacking AAOx production is a consequence of methionine accumulation.

**Significance Statement:** Aliphatic aldoximes are precursors of various specialized metabolites including defense compounds. This study reveals that aliphatic aldoximes repress phenylpropanoid production and that altered methionine metabolism affects plant growth and development. As phenylpropanoids include vital metabolites such as lignin, a major sink of fixed carbon, this metabolic link may contribute to available resource allocation during defense.

## Introduction

Plants produce diverse specialized metabolites that play roles in plant stress adaptation (Pourcel et al., 2007; Chong et al., 2009; Luu et al., 2017; Sørensen et al., 2018; Sugiyama and Hirai, 2019). These specialized metabolites are synthesized through their own biosynthesis pathways. Oftentimes, however, metabolic pathways are interconnected; the alteration of one metabolic pathway can affect the biosynthesis or regulation of other metabolic pathways (Kim et al., 2015; Guo et al., 2016; Mostafa et al., 2016; Nintemann et al., 2017; Xu et al., 2018; Kim et al., 2020; Yang et al., 2020). Analysis of this interconnected nature of plant metabolism is essential for expanding our understanding of how plants coordinate diverse specialized metabolites to adapt to a rapidly changing environment. One example of a metabolic network in specialized metabolism is found in Brassicales, which links together the biosynthesis of glucosinolates and phenylpropanoids (Hemm et al., 2003; Kim et al., 2015; Zhang et al., 2020; Perez et al., 2021). Glucosinolates are Brassicales-specific and structurally diverse defense metabolites (Brader et al., 2006; Halkier and Gershenzon, 2006; Blažević et al., 2020). Glucosinolates are derived from various amino acids; for example, Arabidopsis accumulates glucosinolates derived from tryptophan, phenylalanine and chain-elongated methionine (Kliebenstein et al., 2001; Harun et al., 2020). Phenylpropanoids refer to a class of specialized metabolites mainly derived from phenylalanine and include lignin, flavonoids, and hydroxycinnamates that are crucial for plant growth, defense, and plant-environment interactions (Bonawitz and Chapple, 2010; Pascual et al., 2016; Muro-Villanueva et al., 2019; Dong and Lin, 2021).

While glucosinolates and phenylpropanoids are synthesized through their respective biosynthetic pathways, recent findings have demonstrated that phenylpropanoid biosynthesis can be altered by the glucosinolate intermediate aldoximes (Kim et al., 2015; Kim et al., 2020; Perez et al., 2021). Aldoximes are amino acid derivatives that serve as precursors for various specialized metabolites including glucosinolates and camalexin in Brassicales and cyanogenic glycosides and nitrogenous volatiles throughout the plant kingdom (Glawischnig et al., 2004; Luck et al., 2016; Yamaguchi et al., 2016; Sørensen et al., 2018; Dhandapani et al., 2019). Aldoximes are mainly formed by the action of cytochrome P450 monooxygenases belonging to the 79 family (CYP79) or flavin-containing monooxygenases ((Hansen et al., 2018; Thodberg et al., 2018; Dhandapani et al., 2019; Lai et al., 2020; Liao et al., 2020; Thodberg et al., 2020; Yamaguchi et al., 2021). In Arabidopsis the major aldoxime-forming enzymes are CYP79B2 and CYP79B3 which convert tryptophan to indole-3-acetaldoxime (IAOx) (Hull et al., 2000; Mikkelsen et al., 2000; Zhao et al., 2002); CYP79A2 which converts phenylalanine to phenylacetaldoxime (PAOx) (Wittstock and Halkier, 2000); CYP79F1 and CYP79F2 which generate aliphatic aldoximes (AAOx) from chain-elongated methionine (Chen et al., 2003); and CYP79C1 and CYP79C2 which are promiscuous aldoxime formation enzymes based on a heterologous expression study (Wang et al., 2020) that are barely detected in any organs (Klepikova et al., 2016).

A link among glucosinolates, phenylpropanoids, and aldoximes was first hinted with the isolation of two glucosinolate biosynthesis mutants *reduced epidermal fluorescence 2* (*ref2*) and *ref5* from phenylpropanoid deficient mutant screens (Hemm et al., 2003; Kim et al., 2015). *ref2* mutants contain nonsense mutations in *CYP83A1/REF2* and *ref5* has a missense mutation in *CYP83B1/REF5*. REF2 and REF5 convert aldoximes to their corresponding nitrile oxides in glucosinolate biosynthesis. Although both REF2 and REF5 have affinity for all aldoximes produced in Arabidopsis, these enzymes have different substrate preferences with REF2 having greater affinity for aliphatic aldoximes while REF5 preferentially acts upon aromatic aldoximes (Bak and Feyereisen, 2001; Naur et al., 2003). Consistently, *ref2* mutants accumulate lower levels of aliphatic glucosinolates (Hemm et al., 2003), while *ref5* mutants have reduced indole glucosinolates (Kim et al., 2015). In addition, *ref2* and *ref5* mutants contain reduced levels of phenylpropanoids such as sinapoylmalate, a phenylpropanoid that accumulates in the leaves of Arabidopsis (Hemm et al., 2003; Kim et al., 2015). Further studies have shown that the phenylpropanoid repression in *ref5* is caused not by reduced glucosinolate contents but instead by the accumulation of IAOx or its derivatives since blocking IAOx production rescues phenylpropanoid repression in *ref5* (Kim et al., 2015) and the overproduction of IAOx by *CYP79B2* overexpression reduces phenylpropanoids in Arabidopsis and *Camelina sativa* (Kim et al., 2015; Zhang et al., 2020). Recently it was shown that PAOx accumulation also represses phenylpropanoid production (Perez et al., 2021). One mechanism underlying this glucosinolate-phenylpropanoid crosstalk is increased degradation of phenylalanine ammonia lyase (PAL), the first enzyme of phenylpropanoid biosynthesis, through the transcriptional activation of *Kelch-domain containing F-Box* (*KFB*) genes that target the PAL enzyme for ubiquitination and degradation (Zhang et al., 2013; Zhang and Liu, 2015; Yu et al., 2019; Kim et al., 2020; Perez et al., 2021).

Aliphatic amino acid-derived aldoximes are found widely in the plant kingdom (Sørensen et al., 2018), yet it is unclear if aliphatic aldoximes (AAOx) can repress phenylpropanoid production. Due to high substrate specificity of REF2 toward AAOx (Hemm et al., 2003; Kim et al., 2015; Kim et al., 2020), it was assumed that the reduced phenylpropanoid phenotype of *ref2* is due to the accumulation of AAOx (Hemm et al., 2003). However, given the range of substrates for REF2, it is possible that other aldoximes such as IAOx may fully or partially contribute to phenylpropanoid repression in *ref2*. Indeed, Bak and Feyereisen (2001) demonstrated that *REF2* overexpression can rescue the high auxin phenotype of the *REF5* null mutant *rnt1-1*, implying that REF2 can act upon IAOx *in vivo*. IAOx and PAOx are single molecules derived from tryptophan and phenylalanine respectively and they are precursors of the natural auxins IAA and PAA respectively (Zhao et al., 2002; Perez et al., 2021; Perez et al., 2022). On the other hand, AAOx structure varies depending on the length of chain-elongated methionine. In Arabidopsis, CYP79F1 and CYP79F2 produce all AAOx from diverse chain-elongated methione (Chen et al., 2003; Tantikanjana et al., 2004). The impact of AAOx on phenylpropanoid production can be tested through the overexpression of *CYP79F1*/*F2* or by removing CYP79F1/F2 activities. However, several studies have shown that overexpression of either *CYP79F1* or *CYP79F2* co-suppresses both *CYP79F1* and *CYP79F2*, which leads to reduced rather than increased CYP79F1/F2 activities (Hansen et al., 2001; Reintanz et al., 2001; Chen et al., 2003). On the other hand, elimination of AAOx production by knockout or silencing of *CYP79F1* and *CYP79F2* results in severe growth defects such as cup-shaped rosette leaves, loss of apical dominance and sterility (Hansen et al., 2001; Reintanz et al., 2001; Tantikanjana et al., 2001; Chen et al., 2003; Tantikanjana et al., 2004; Chen et al., 2012). Foremost, *CYP79F1* (At1g16410) and *CYP79F2* (At1g16400) are physically linked. While several hypotheses have been made attempting to explain how this altered growth and development come about, including ones which attribute the bushy phenotype to perturbed auxin or cytokinin homeostasis, the mechanism behind this growth phenotype remains elusive (Reintanz et al., 2001; Tantikanjana et al., 2004; Chen et al., 2012).

In this study, we examined the impact of AAOx metabolism on phenylpropanoid production in Arabidopsis using various glucosinolate biosynthesis mutants. Additionally, upstream elements of AAOx production were examined to further elucidate the mechanism of altered growth in plants lacking AAOx production. Our metabolic profiling and genetic study demonstrated the impact of altered AAOx metabolism on phenylpropanoid production and revealed that the abnormal developmental phenotypes of *CYP79F1/CYP79F2* silenced plants is due to a disruption in methionine homeostasis.

## Results

### IAOx Accumulates in *ref2*

REF2 has high affinity for AAOx while REF5 has greater affinity for IAOx than REF2 (Figure 1a) (Bak and Feyereisen, 2001; Naur et al., 2003). Consistently, *ref5* mutants have decreased indole glucosinolates and display high auxin morphological phenotypes due to increased IAA which is redirected from accumulated IAOx (Delarue et al., 1998; Barlier et al., 2000; Bak and Feyereisen, 2001; Hoecker et al., 2004). On the other hand, *ref2* mutants have decreased aliphatic glucosinolates and look similar to wild type (Hemm et al., 2003; Kim et al., 2015; Kim et al., 2020). Surprisingly, we detected a significant level of IAOx in the *ref2* mutant (Figure 1b, 1c). This result is unexpected because REF5, the major IAOx catabolic enzyme, is still functional in *ref2* and the *ref2* mutant does not display high auxin phenotypes which was observed in other IAOx accumulation mutants such as *ref5* or *sur1* (Delarue et al., 1998; Barlier et al., 2000; Mikkelsen et al., 2004; Kim et al., 2015). As IAOx accumulation represses phenylpropanoid biosynthesis (Kim et al., 2015; Zhang et al., 2020), all or part of the reduction in phenylpropanoid content seen in *ref2* is likely due to this IAOx accumulation, which leaves the question whether AAOx has any impact on phenylpropanoid metabolism.

**Figure 1.**
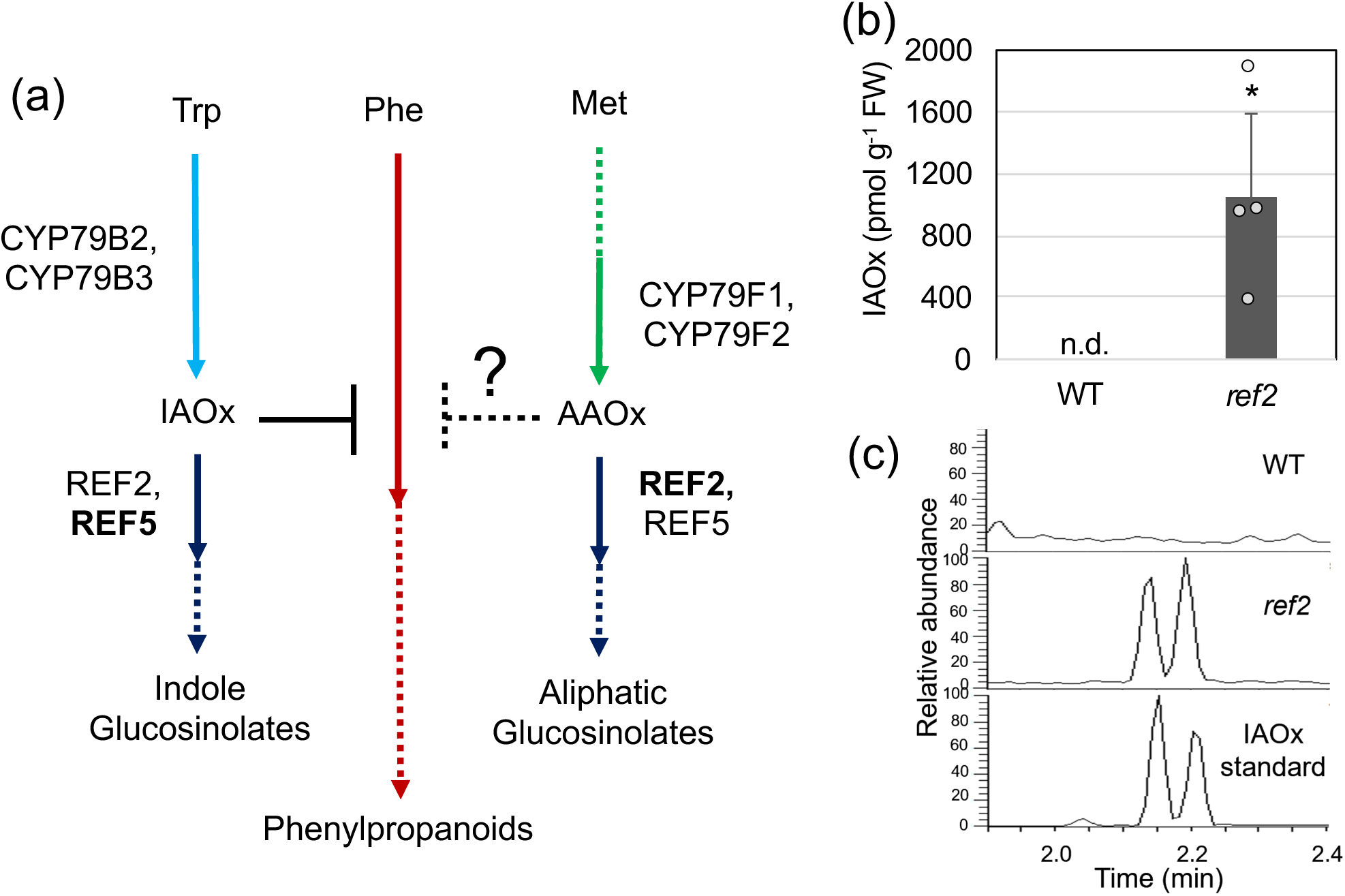
A metabolic link between aldoxime metabolism and the phenylpropanoid pathway in Arabidopsis. (a) A schematic diagram of a metabolic link between the phenylpropanoid pathway and glucosinolate production in *Arabidopsis thaliana*. It was previously shown that the accumulation of indole 3-acetaldoxime (IAOx), an indole glucosinolate intermediate, represses the phenylpropanoid pathway. The impact of aliphatic aldoximes (AAOx) on the phenylpropanoid pathway remains unknown. Although REF2 and REF5 function redundantly to metabolize all aldoximes, REF2 is the preferred enzyme for converting AAOx. (b) IAOx content in leaves of two-week-old wild type and *ref2.* Data represents mean ± SD (n=4). The individual data points are shown on the bar graphs. The means were compared by Student’s t-test and statistically significant differences (*P*<0.05) are indicated by an asterisk (*). (c) LC-MS chromatograms show accumulation of IAOx in *ref2*. *E*-IAOx and *Z*-IAOx standards (bottom) and extracts from wild type (top) and *ref2* (middle) are shown.

To examine the impact of AAOx on phenylpropanoid production, we first tested if accumulated IAOx in *ref2* is entirely responsible for phenylpropanoid repression by removing IAOx production in *ref2*. As *cyp79b2 cyp79b3* (*b2b3*) double mutants do not produce IAOx (Zhao et al., 2002), *ref2* and *b2b3* were crossed to generate the *b2b3ref2* triple mutant (Figure 2a). As expected, the *b2b3ref2* triple mutant was unable to generate IAOx and IAOx-derived glucosinolates (I3M) while it still produces aliphatic glucosinolates (Figure 2b-d). The level of sinapoylmalate was increased significantly in the *b2b3ref2* compared to *ref2* but was not returned to wild-type levels (Figure 2e). These results confirm an inhibitory effect of IAOx accumulation on phenylpropanoid production but suggest that IAOx accumulation only partially contributes to the repression of phenylpropanoid production in *ref2*.

**Figure 2.**
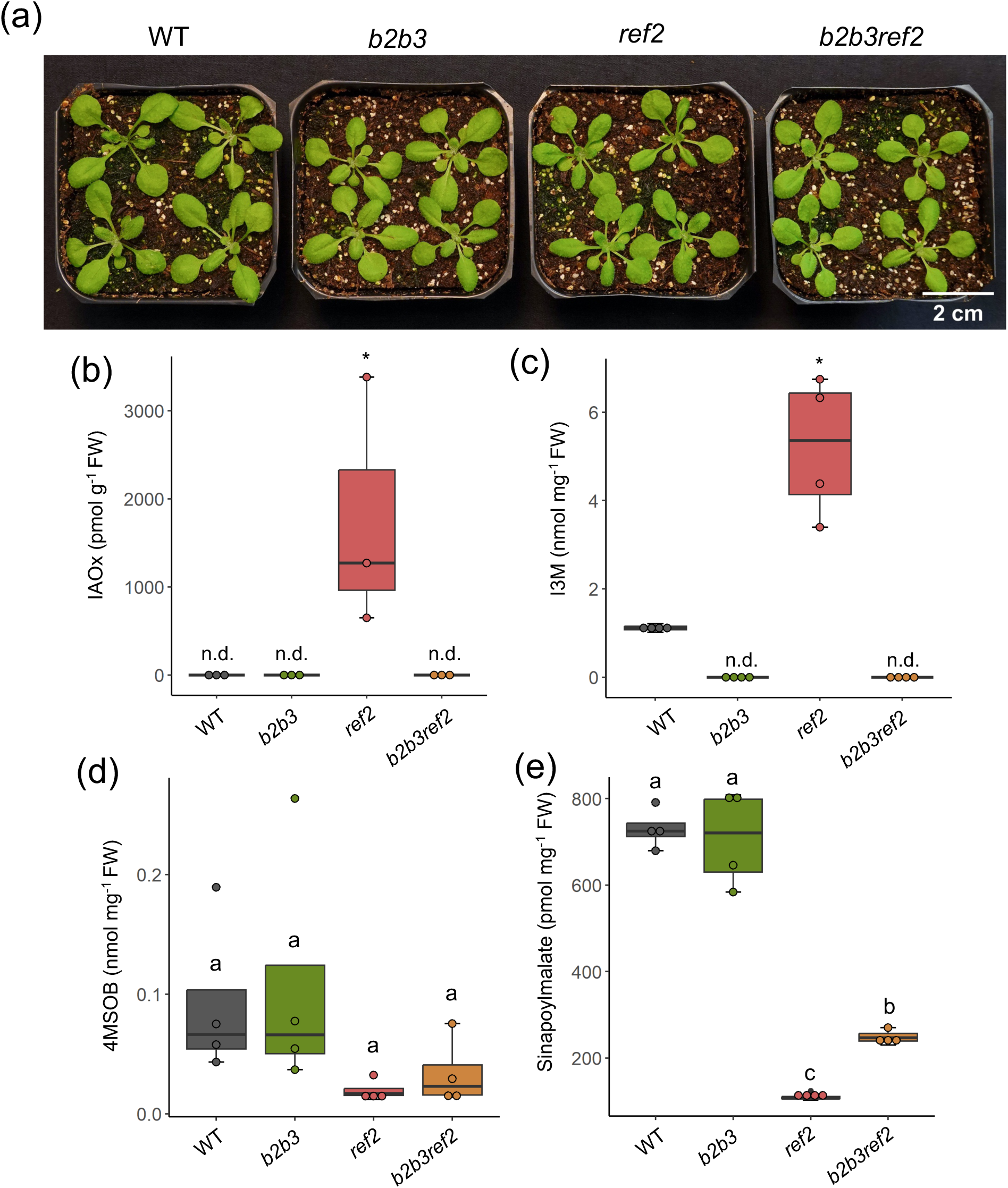
Elimination of IAOx biosynthesis partially rescues phenylpropanoid production in *ref2*. (a) Representative images of three-week-old wild type (WT), *b2b3 (cyp79b2 cyp79b3)* double mutant, *ref2*, and *b2b3ref2 (cyp79b2 cyp79b3 ref2)* triple mutant. (b-e) The levels of indole-3-acetaldoxime (IAOx) (b), desulfo-I3M (c), desulfo-4-methylsulfinylbutyl glucosinolate (4MSOB) (d), and sinapoylmalate (e) were measured from the leaves of three-week-old wild type, *b2b3*, *ref2*, and *b2b3ref2* plants. The box represents the interquartile range, spanning from the first quartile to the third quartile, with the median depicted by a line within the box. Whiskers extend to the minimum and maximum values. The individual data points are displayed on the bar graphs. Student’s t-test was used to compare the means of IAOx and I3M, and statistically significant differences (P<0.05) are denoted by an asterisk (*) (n=3 for IAOx, n=4 for I3M). The statistical significance of 4MSOB and sinapoylmalate contents was determined using a one-way ANOVA, with a significance level set at P<0.05. Differences among groups were further analyzed using Tukey’s post-hoc test, and significant differences (n=4) are indicated by letters. ‘n.d.’ is not detected.

### AAOx Accumulation Represses Phenylpropanoid Production

The metabolic profile of the *b2b3ref2* triple mutant (Figure 2) implies that another mechanism besides IAOx accumulation in *ref2* results in phenylpropanoid repression. As REF2 is considered the major enzyme for aliphatic glucosinolate biosynthesis, we hypothesized that *ref2* accumulates AAOx, which represses phenylpropanoid production. To test our hypothesis, we chose to reduce all AAOx production in *ref2* by disrupting both CYP79F1 and CYP79F2 simultaneously. Due to the tandem position of the *CYP79F1* (*At1g16410*) and *CYP79F2* (*At1g16400*) genes, removal of AAOx by generating a *cyp79f1 cyp79f2* double mutant is challenging. However, several studies have shown that overexpression of either *CYP79F1* or *CYP79F2* using a strong promoter results in cosuppression of both *CYP79F1* and *CYP79F2*, which leads to display characteristic *cyp79f1* or *cyp79f2* loss-of function mutant phenotypes such as curled-up leaf morphology and bushy stems (Hansen et al., 2001; Reintanz et al., 2001; Chen et al., 2003; Tantikanjana et al., 2004). Thus, we decided to remove AAOx production via expressing *CYP79F1* using the CaMV35S promoter in wild type and *ref2* and examined how silencing of *CYP79F1* and *CYP79F2* affects phenylpropanoid production.

A majority of T1 transgenic plants displayed the atypical growth and developmental phenotypes associated with *CYP79F1/CYP79F2* co-suppression (Figure 3a), which is consistent with previous reports (Hansen et al., 2001; Reintanz et al., 2001; Chen et al., 2003; Tantikanjana et al., 2004). The rosette leaves were curled-up (or cup-shaped) (Figure 3a) and mature plants produced multiple stems leading to a “bushy” phenotype (Figure 3a); most plants did not produce seeds under our growth conditions. As reported previously, *CYP79F1* and *CYP79F2* expression was found to be reduced in the “bushy” transgenic lines compared to their controls (Figure 3a). We named these “bushy” *CYP79F1*/*CYP79F2* cosuppression plants *F1-cos/WT* and *F1-cos/ref2,* which are *F1-*cosuppression lines in wild type and *ref2* backgrounds respectively. About ten percent of T1 transgenic plants did not display the characteristic “bushy” and “cup-shaped” phenotype (Figure 3a), which is consistent with a previous study (Hansen et al., 2001; Reintanz et al., 2001; Chen et al., 2003). Thus, we named these plants *F1-OX/WT* and *F1-OX/ref2*.

**Figure 3.**
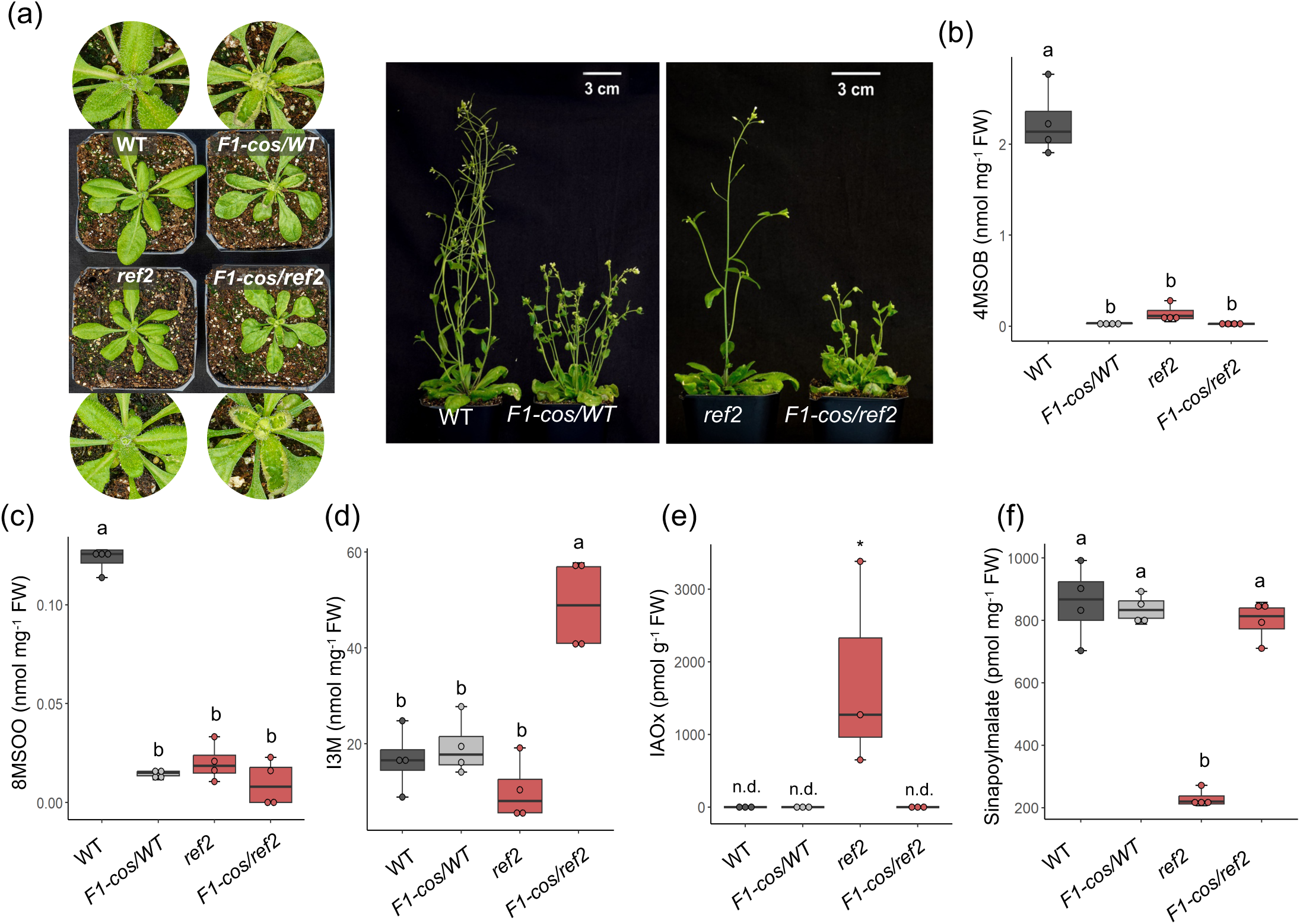
*CYP79F1*-cosuppression restores phenylpropanoid contents in *ref2*. (a) Representative images of three-week-old and six-week-old plants. *F1-cos* lines exhibit distinctive morphological phenotypes such as curled-up leaves and “bushy”-like stems. (b-f) The levels of desulfo-4-methylsulfinylbutyl glucosinolate (4MSOB) (b), desulfo-8-methylsulfinyloctyl glucosinolate (8MSOO) (c), desulfo-I3M (d), indole-3-acetaldoxime (IAOx) (e) and sinapoylmalate (f) of wild type, *F1*-cosuppression lines in four-week-old wild type (*F1-cos/WT*), *ref2*, and *F1-cos* lines in *ref2* (*F1-cos/ref2*). The box represents the interquartile range, spanning from the first quartile to the third quartile, with the median indicated by a line within the box. Whiskers extend to the minimum and maximum values. The individual data points are displayed on the bar graphs. The statistical significance of the glucosinolate and sinapoylmalate results was determined using a one-way ANOVA, with a significance level set at P<0.05 (n=4). Differences among groups were further analyzed using Tukey’s post-hoc test and significant differences are indicated by letters. Student’s t-test was used to compare the means of IAOx and statistically significant differences (P<0.05) are denoted by an asterisk (*) (n=3). ‘n.d.’ is not detected.

To confirm disruption of CYP79F1 and CYP79F2 activities in *F1-cos* lines, the glucosinolate profiles of mature T1 *F1-cos* lines in the wild type (*F1-cos/WT*) and *ref2* genetic backgrounds (*F1-cos/ref2)* were determined. In *F1-cos* lines of both genetic backgrounds, production of short-chain and long-chain AAOx-derived glucosinolates (4-methylsulfinylbutyl glucosinolate [4MSOB] and 8-methylsulfinyloctyl glucosinolate [8MSOO] respectively) was almost completely eliminated while it accumulates indole glucosinolates, indicating the lack of CYP79F1 and CYP79F2 activities (Figure 3b-d). Although the levels of aliphatic glucosinolates (4MSOB and 8MSOO) in *F1-cos/ref2* are comparable to that in *ref2, ref2* has decreased aliphatic glucosinolates because of its defect in the conversion of AAOx to their respective nitrile oxides whereas decreased aliphatic glucosinolates in *F1-cos/ref2* plants are due to a reduction in AAOx production. Interestingly, IAOx content in *F1-cos/ref2* is under the detection limit (Figure 3e). We then compared phenylpropanoid contents in *ref2* and *F1-cos/ref2* and found that *F1-cos/ref2* lines produce wild-type levels of sinapoylmalate (Figure 3f). The fact that disruption of CYP79F1/F2 restored phenylpropanoid production in *ref2* suggests that AAOx accumulation exerts a repression effect on phenylpropanoid production.

Regarding IAOx-mediated and PAOx-mediated phenylpropanoid repression, the mechanism underlying this metabolic interaction includes increased PAL degradation, which ultimately reduces PAL activity and total phenylpropanoid production (Kim et al., 2020; Zhang et al., 2020; Perez et al., 2021). To test if F1-cosuppression has any impact on PAL activity, we measured PAL activity in *F1-cos* lines. In line with previous reports, *ref2* showed reduced PAL activity compared to wild type (Figure 4). In *F1-cos/ref2* lines, however, PAL activity was restored to the wild-type level (Figure 4). Ultimately these results demonstrate that depletion of AAOx production restores decreased PAL activity in *ref2*.

**Figure 4.**
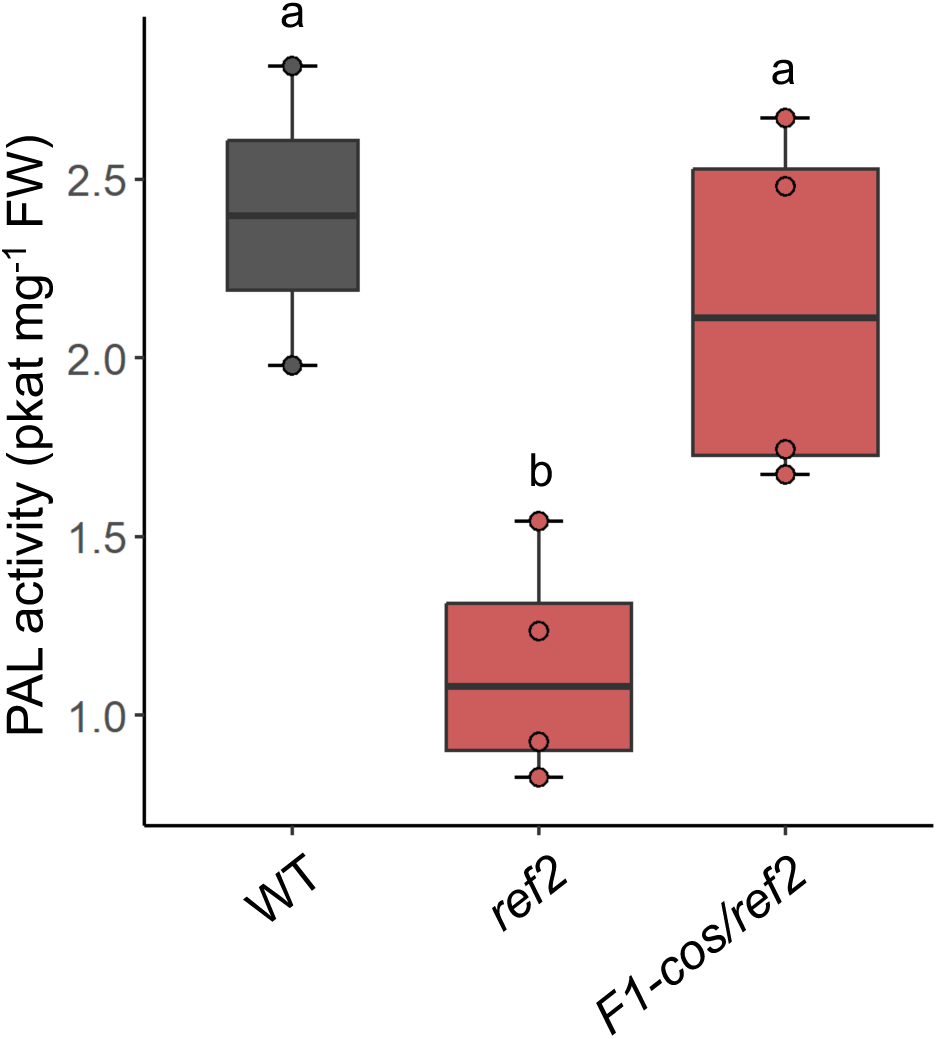
PAL activity is restored in *F1-cos/ref2*. PAL activity of wild type, *ref2* and *F1-cos/ref2*. PAL activities were measured using leaves from four-week-old soil grown plants. The box represents the interquartile range, spanning from the first quartile to the third quartile, with the median indicated by a line within the box. Whiskers extend to the minimum and maximum values. The individual data points are shown on the graphs. The statistical significance of the results was determined through a one-way ANOVA, with a significance level set at P<0.05. Differences among groups were further analyzed using Tukey’s post-hoc test and significant differences (n=2 for wild type, n=4 for *ref2* and *F1-cos/ref2*) are indicated by letters.

We also generated *F1-cos* lines in *ref5* background (*F1-cos/ref5*). They consistently displayed characteristic abnormal morphological phenotypes resulting from *CYP79F1/F2* cosuppression and contained significantly reduced aliphatic glucosinolate contents and increased indole glucosinolate contents compared to wild type and *ref5* (Figure 5a-d). Notably, *F1-cos/ref5* accumulates more sinapoylmalate than *ref5,* suggesting redundant function of REF5 in AAOx conversion (Figure 5e).

**Figure 5.**
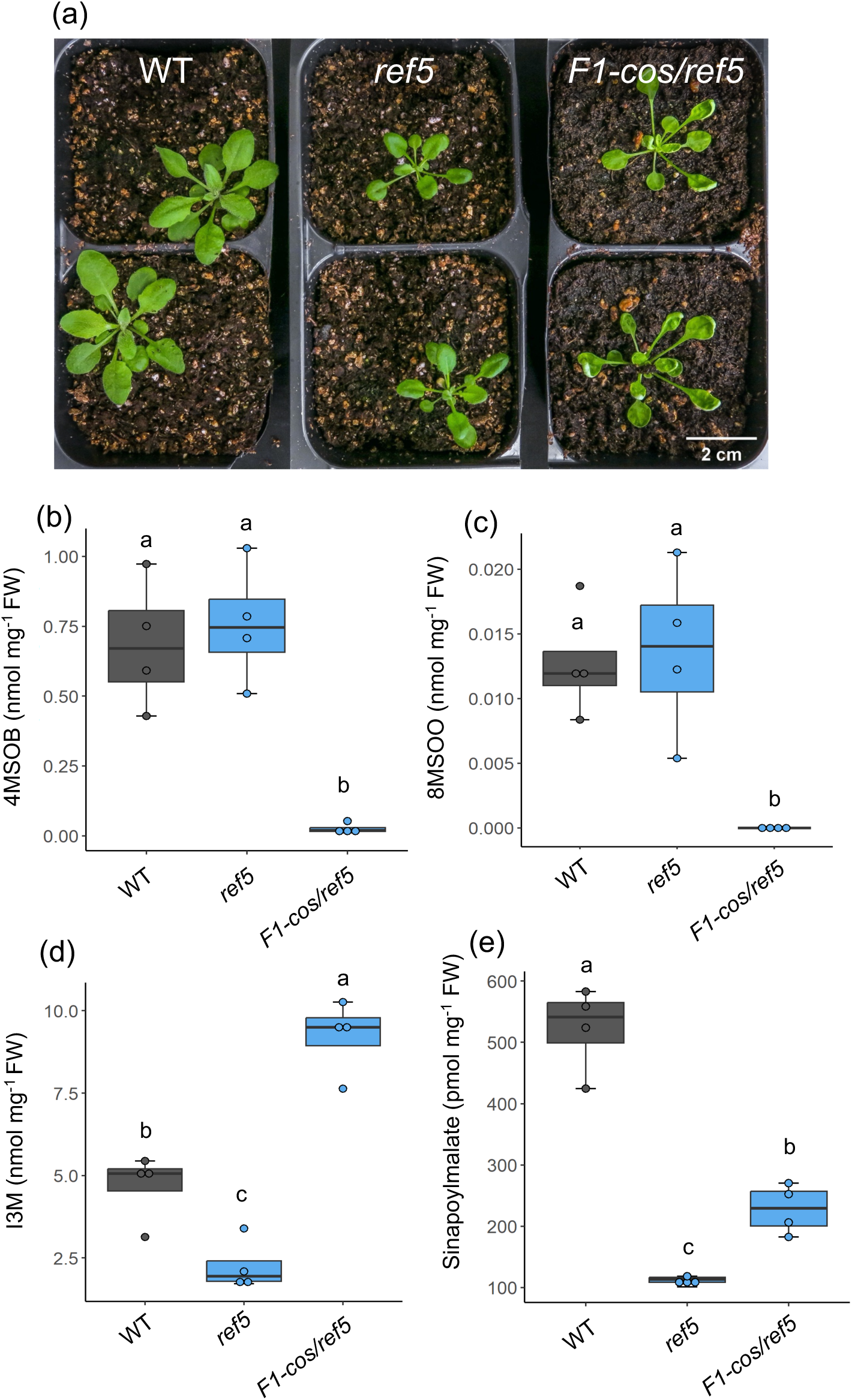
Cosuppression of *CYP79F1* and *CYP79F2* partially rescue phenylpropanoid contents in *ref5*. (a) Representative image of three-week-old wild type, *ref5,* and *F1-cos* lines in the *ref5* genetic background (*F1-cos/ref5)*. (b-e) The levels of desulfo-4-methylsulfinylbutyl glucosinolate (4MSOB) (b), desulfo-8-methylsulfinyloctyl glucosinolate (8MSOO) (c), desulfo-I3M (d) and sinapoylmalate (e) of wild type, *ref5*, and *F1-cos/ref5*. The box represents the interquartile, spanning from the first quartile to the third quartile, with the median indicated by a line within the box. Whiskers extend to the minimum and maximum values. The individual data points are shown on the graphs. The statistical significance of the results was determined through a one-way ANOVA, with a significance level set at P<0.05. Differences among groups were further analyzed using Tukey’s post-hoc test and significant differences (n=4) are indicated by letters.

### IAA Content is Unaffected in *F1-cos* Lines

Although the unique growth and developmental changes of *CYP79F1/CYP79F2* knockout or silenced plants were observed in several studies, how the removal of CYP79F1 or CYP79F2 activity results in unique growth alteration remains unanswered. It was proposed that misregulation of auxin homeostasis may play a role in this alteration (Reintanz et al., 2001; Tantikanjana et al., 2001). However, a more recent report by Chen et al. (2012) found that IAA levels were unchanged or slightly reduced in *CYP79F1* RNAi lines.

To determine how *CYP79F1/CYP79F2* cosuppression impacts IAA biosynthesis, we quantified IAA content in wild-type, *ref2* and *F1-cos/ref2* plants. Despite accumulating significant levels of IAOx (Figure 1b), *ref2* plants were found to contain wild-type levels of IAA (Figure 6), an observation which explains why *ref2* mutants do not display a high auxin growth morphology. Similar to *ref2* plants, IAA content in the leaves of 3-week-old *F1-cos/ref2* lines was not different from that in wild type under our growth conditions (Figure 6). It is noteworthy that *ref5* contains increased auxin and displays high auxin morphology including hyponasty (curled-down) leaf morphology (Kim et al., 2015), but *F1-cos/ref5* showed curled-up leaf morphology (Figure 5a). It is unlikely that the growth phenotype of our *F1-cos* line or of *cyp79f1* and *cyp79f2* mutants is the result of abnormal auxin metabolism.

**Figure 6.**
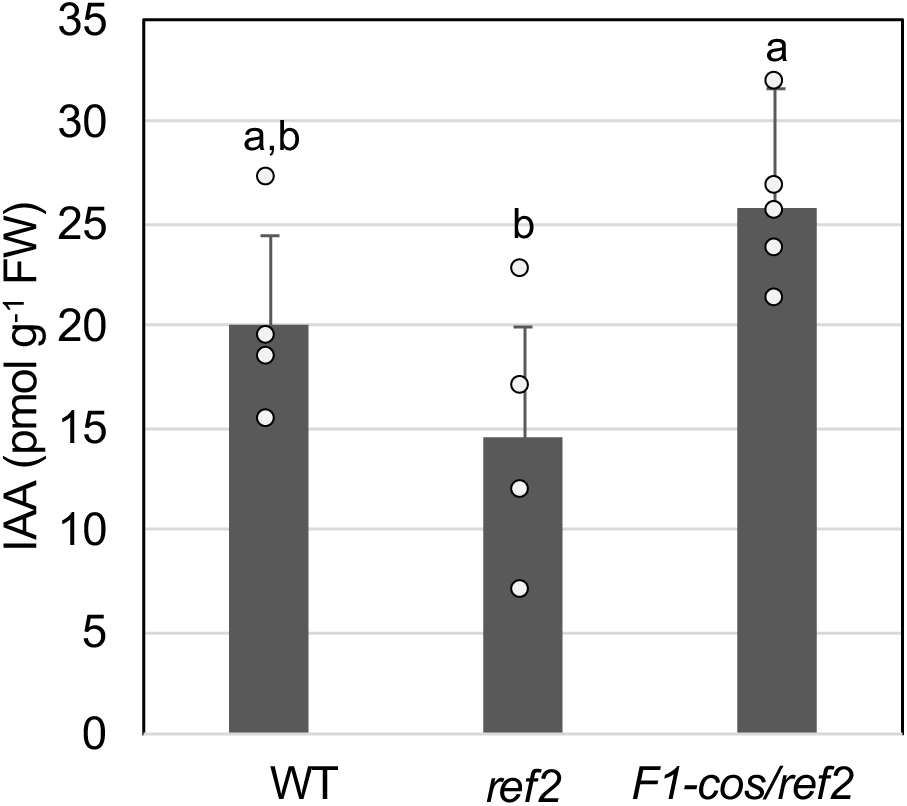
IAA content is unaltered in *F1-cos/ref2* plants compared to wild type and *ref2*. Free IAA content of three-week-old wild type, *ref2*, and *F1-cos/ref2* plants. Data represents mean ± SD (n=4). The individual data points are shown on the graph. The statistical significance of the results was determined through a one-way ANOVA. Differences among groups were further analyzed using Tukey’s post-hoc test (P<0.05) and were identified by letters to indicate significance.

### Methionine Feeding Phenocopies Growth Morphology of *F1-cos* Lines

It was previously shown that methionine content is increased in some aliphatic glucosinolate-deficient mutants including *CYP79F1*-silenced plants (Sawada et al., 2009; Chen et al., 2012). Consistently, *F1-cos/ref2* contains increased methionine compared to *ref2* (Figure 7). Since *ref2* mutants downstream of CYP79F1/F2 do not show the altered growth phenotypes observed in *cyp79f1/cyp79f2* mutants or *F1-cos* lines, we hypothesized that the accumulation of methionine or a methionine-derived metabolite (barring AAOx) might be responsible for the altered growth of our *F1-cos* lines and other *cyp79f1/cyp79f2* mutants.

**Figure 7.**
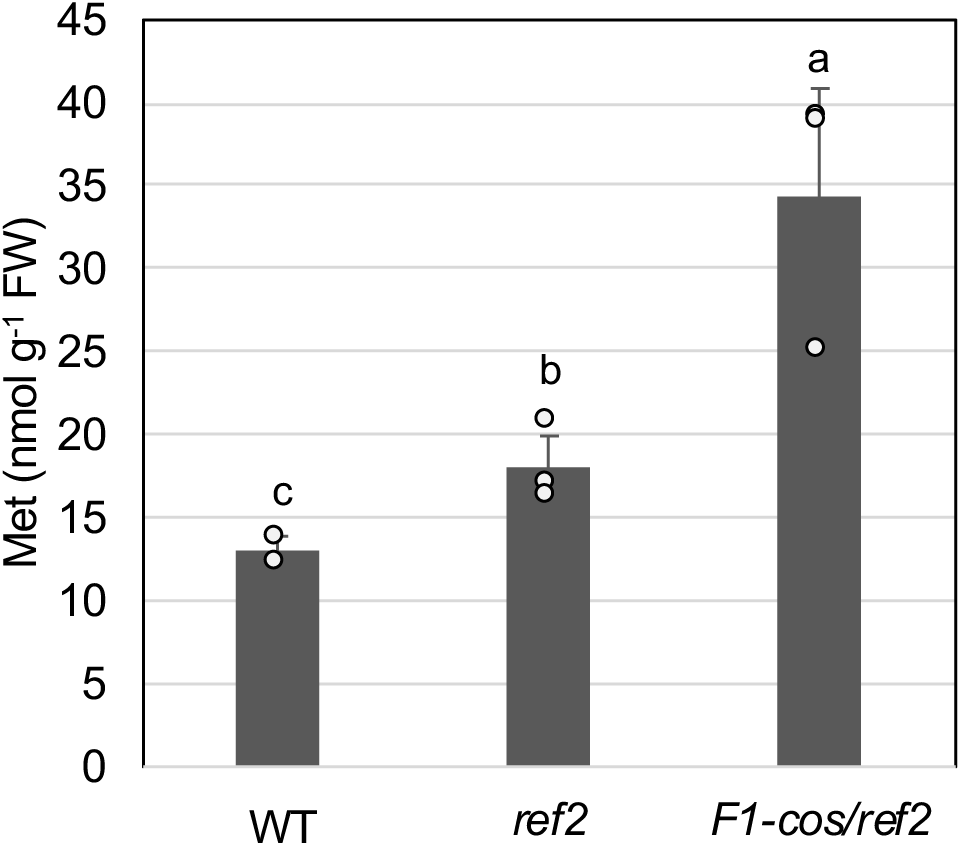
**The methionine content in *F1-cos/ref2* is higher than in wild type and *ref2***. Methionine contents were measured with three-week-old wild type, *ref2*, and *F1-cos/ref2* plants. Data represents mean ± SD (n=3). The individual data points are shown on the bar graph. The statistical significance of the results was determined through a one-way ANOVA, with a level of significance set at P<0.05. Differences among groups were further analyzed using Tukey’s post-hoc test and were identified by letters to indicate significance.

To examine any impact of methionine accumulation on plants growth, wild type and *ref2* plants were grown on growth media with or without methionine. After three weeks, plants of both genetic backgrounds grown on plates supplemented with 300 μM of methionine began displaying the cup-shaped leaf morphology associated with loss of CYP79F1/CYP79F2 activity (Figure 8a, 8b). While control plants and methionine-treated plants accumulated similar levels of sinapoylmalate (Figure 8c), methionine-treated wild-type plants contained increased aliphatic glucosinolates, but not indole glucosinolates (Figure 8d-f). S-adenosyl methionine (SAM) is a methionine derivative and a precursor of both the plant hormone ethylene and polyamines (Figure 9). To determine if the accumulation of these metabolites causes the altered growth morphology, we fed three polyamines (putrescine, spermidine, and spermine) and the ethylene precursor ACC (1-aminocyclopropane-1-carboxylate) to wild type and *ref2* plants, but none of them phenocopied *F1-cos* lines (Figure S2).

**Figure 8.**
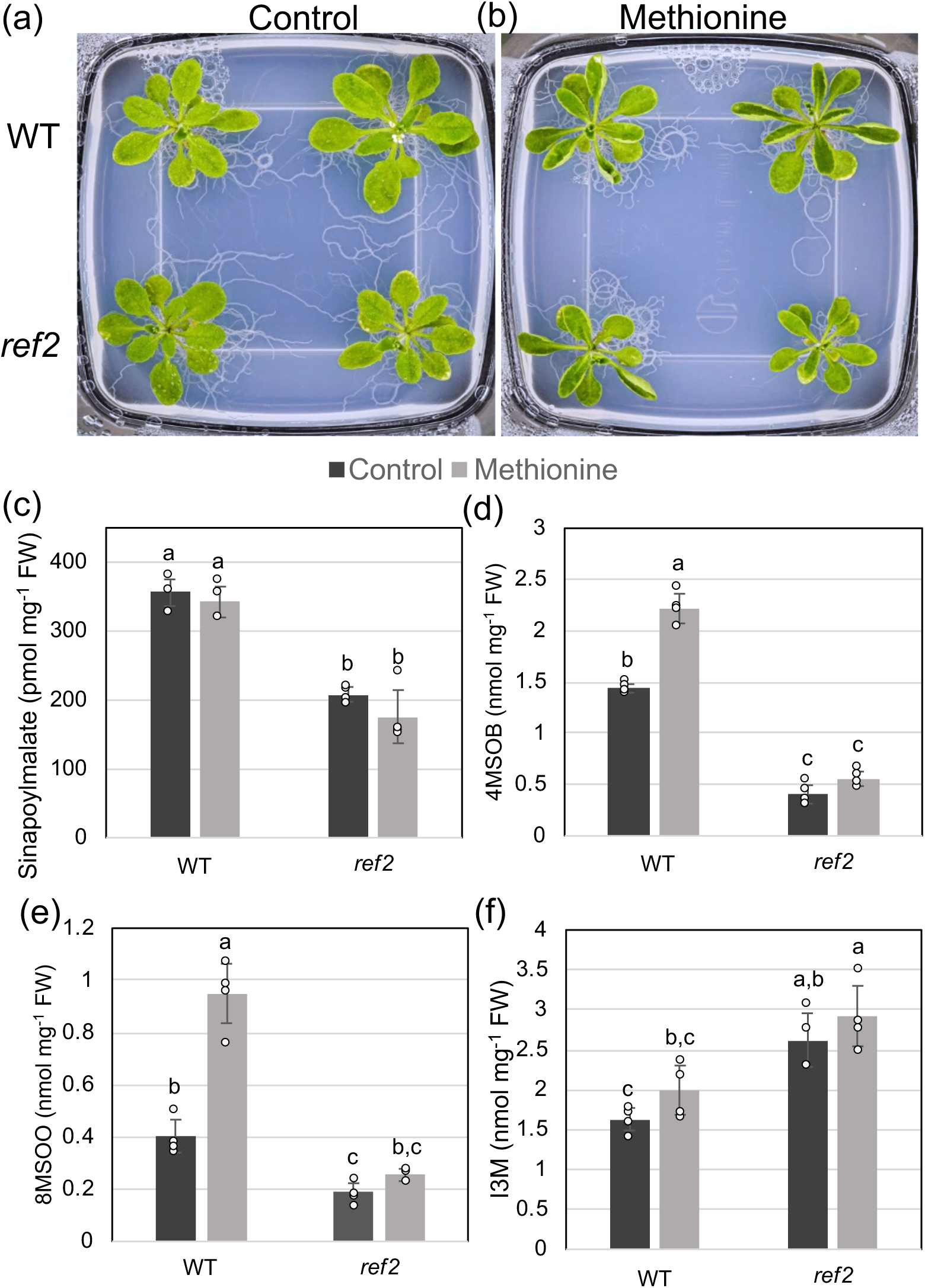
Wild type and *ref2* grown on methionine-containing media show curled-up leaf morphology similar to those shown in *F1-cos* lines. (a-b) Representative images of three-week-old wild type and *ref2* plants grown on MS plates (a) and MS plates supplemented with 300 μM methionine (b). (c-f) The levels of sinapoylmalate (c), desulfo-4MSOB (d), desulfo-8MSOO (e), and desulfo-I3M (f) of three-week-old wild type and *ref2* grown on MS or methionine-supplemented MS plates. Data represents mean ± SD (n=4). The individual data points are shown on the graph. The statistical significance of the results was determined through a one-way ANOVA, with a level of significance set at P<0.05. Differences among groups were further analyzed using Tukey’s post-hoc test and were identified by letters to indicate significance.

**Figure 9.**
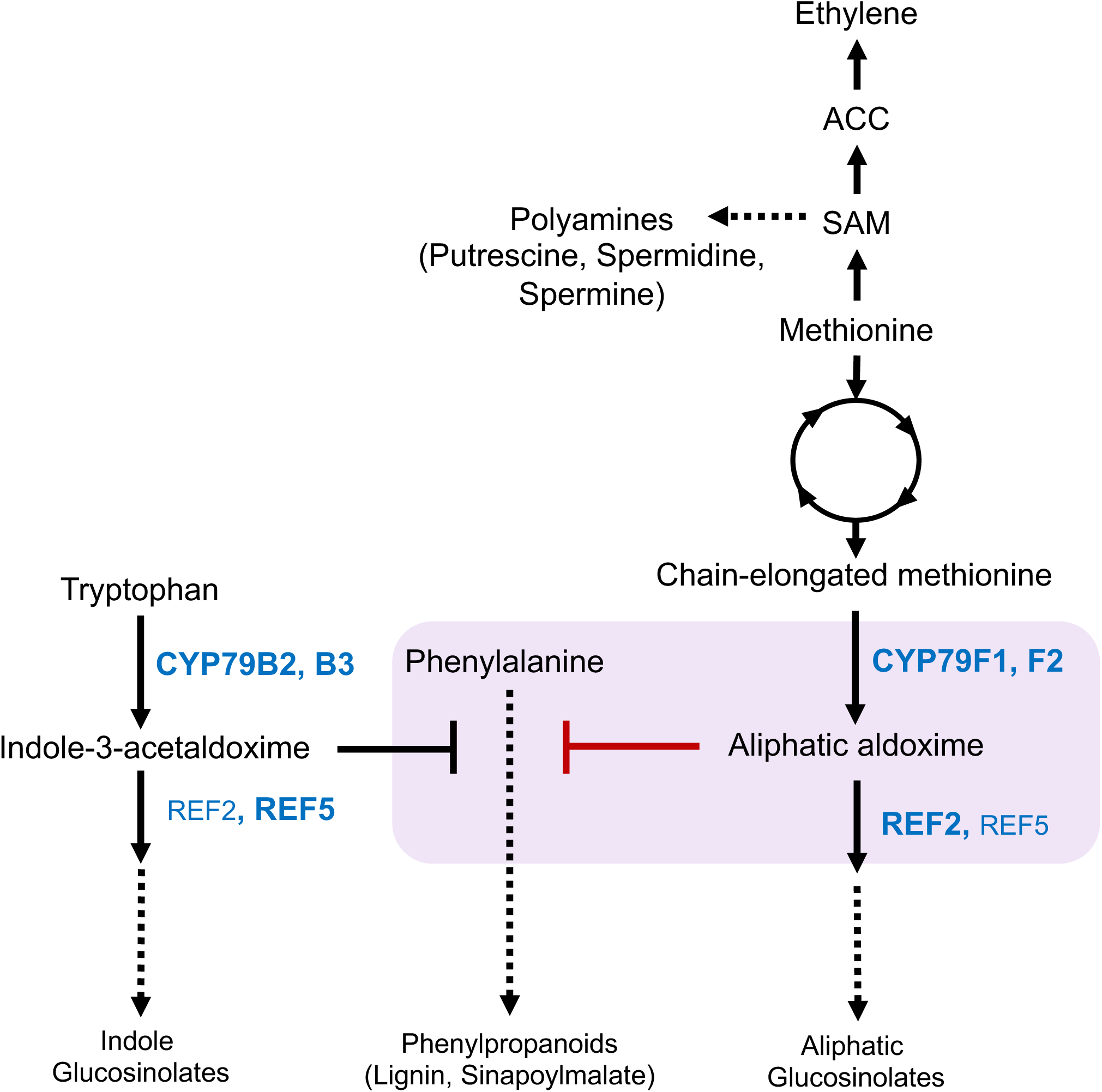
Working model of the impact of methionine-derived aliphatic aldoximes (AAOx) on phenylpropanoid production. A schematic detailing the metabolic interaction between methionine metabolism, glucosinolate biosynthesis, and phenylpropanoid production. Known inhibitory interaction of indole 3-acetaldoxime (IAOx) with the phenylpropanoid pathway is indicated. Production of aliphatic glucosinolates from methionine requires the biosynthesis of AAOx, which upon their accumulation represses phenylpropanoid production (red line in light purple box). Primary enzymes for indole versus aliphatic glucosinolate biosynthesis are bolded. This metabolic interconnection, along with other methionine-derived metabolites such as ethylene and polyamines, can influence plant growth by altering leaf morphology and axillary bud initiation or apical dominance. The metabolic intermediates of the methionine chain elongation pathway may also impact plant growth.

Taken together, our results suggest that the accumulation of AAOx or its derivatives specifically affects phenylpropanoid metabolism partially through repression of PAL activity, while the accumulation of methionine or a non-aldoxime derivative exerts additional effects on plant growth that ultimately results in the curled-up leaf morphology and bushy phenotype.

## Discussion

In this study, we identified a metabolic network whereby methionine and methionine-derived AAOx were shown to significantly impact phenylpropanoid production and overall plant growth (Figure 9). It was demonstrated previously that IAOx-mediated phenylpropanoid repression is the result of the accumulation of IAOx or its derivatives but not IAA or indolic glucosinolates (Kim et al., 2015). Partial restoration of sinapoylmalate content in *ref2* through blockage of IAOx production further confirms the repressive effect of IAOx on phenylpropanoid production (Figure 2). Similarly, the accumulation of AAOx represses phenylpropanoid production, as disruption of *CYP79F1/CYP79F2* increases levels of sinapoylmalate in both *ref2* and *ref5* (Figure 3, 5). Apparently, phenylpropanoid repression in *ref2* and *ref5* appears to be a consequence of both IAOx and AAOx accumulation (Figure 2, 3, 5)(Kim et al., 2015). A recent work has shown that PAOx accumulation also represses phenylpropanoid production (Perez et al., 2021). Given the similar structures of IAOx and PAOx, their repressive effect is not surprising. Unlike IAOx and PAOx, though, AAOx refers to several aldoximes derived from various chain-elongated methionine derivatives including 5-methylthiopentanaldoxime, 6-methylthiohexanaldoxime, and 4-methylthiobutyraldoxime (Matsuo, 1968; Hansen et al., 2001; Tantikanjana et al., 2004). It remains unknown how these and other aldoximes aldoximes are sensed and how these signals are transduced to alter phenylpropanoid metabolism. Nevertheless, it seems that alternative aldoxime-derived compounds or perhaps aldoximes themselves serve as signals to alter phenylpropanoid production. Aliphatic glucosinolates make up a significant portion of the Arabidopsis glucosinolate profile in many tissues including inflorescences (Brown et al., 2003). Various aliphatic aldoximes derived from valine, isoleucine, and leucine are found widely in plants and their production increases under biotic stresses (Sørensen et al., 2018). Given that the altered AAOx production affects the phenylpropanoid pathway which produces an array of specialized metabolites, including lignin, a major sink of fixed carbon, this metabolic link may contribute to the allocation of available resources under stress.

The discovery of IAOx accumulation in *ref2* (Figure 1b, 1c) is surprising in and of itself as *ref2* does not accumulate IAA in the same way as removal of REF5 activity, where IAOx accumulation results in increased IAA production (Delarue et al., 1998; Barlier et al., 2000; Bak and Feyereisen, 2001; Hoecker et al., 2004). Since *ref2* mutants contain higher levels of indolic glucosinolates compared to wild-type plants (Figure 2b) (Hemm et al., 2003), this IAOx accumulation is likely from increased flux towards indole glucosinolates in *ref2*. It remains unclear why the excess IAOx in *ref2* does not lead to increase IAA.

In terms of the unique growth morphology of *F1-cos* lines, several studies suggest that alterations in auxin and cytokinin homeostasis may be the causal agent as increased levels of IAA and cytokinins have been detected in *F1-cos* lines previously (Tantikanjana et al., 2001; Tantikanjana et al., 2004). On the other hand, Chen et al. (2012) demonstrated that free auxin was unchanged in *CYP79F1*-silenced plants, and a similar result was observed in this study (Figure 6). Discrepancy of auxin contents might be from sample variation or growth conditions, given pleotropic morphological changes in *CYP79F1-cos* lines. Arabidopsis plants having increased auxins such as YUC overexpression lines and *ref5* often display epinasty leaf morphology (curled-down)(Zhao et al., 2001; Kim et al., 2007; Kim et al., 2015). It is noteworthy that cosuppression of *F1/F2* reverses epinasty leaves of *ref5* to be curled-up (Figure 5a). Nonetheless, auxin is unlikely the main player in these developmental changes particularly the cup-shaped leaf morphology in *CYP79F1-cos* lines. We showed that an increase in methionine or methionine-derived metabolite(s) results in cup-shaped leaves in both the wild type and *ref2* genetic backgrounds (Figure 8), suggesting that the metabolite responsible for this morphology is upstream of or unrelated to aliphatic glucosinolate biosynthesis. Methionine is an essential amino acid that serves as a precursor of *S*-adenosyl-methionine, which serves as both a major methyl donor as well as an intermediate in the biosynthesis of the phytohormones ethylene and polyamines (Wang et al., 2002; Handa et al., 2018). It is therefore possible that elimination of AAOx production in Arabidopsis results in accumulation of methionine which in turn affects flux towards the biosynthesis of these methionine-derived metabolites. Although none of the plants grown on growth media supplemented with polyamines and the ethylene precursor ACC phenocopied *F1-cos* lines under our growth conditions (Figure S2), further study is necessary to rule out the effect of any hormonal interactions on the characteristic growth alteration. It is also possible that the *F1-cos* growth phenotype may be more generally the result of misregulated amino acid metabolism, as it was recently shown that feeding of Arabidopsis with tyrosine causes cup-shaped leaf morphology (Yokoyama et al., 2021).

In summary, this study demonstrated that primary and specialzed metabolism related to methionine can significantly impact plant growth in Arabidopsis through AAOx-mediated phenylpropanoid repression as well as through other processes or metabolites. While these results have currently only been uncovered in Arabidopsis, methionine and phenylpropanoid metabolism are universally distributed throughout the plant kingdom, and regulatory mechanisms governing activity or flux through these pathways are oftentimes conserved among many species. Additionally, aldoxime metabolism and particularly AAOx production is widespread throughout the plant kingdom, and recent findings demonstrating the conservation of other aspects of aldoxime metabolism such as auxin production outside of the Brassicales family (Perez et al., 2021; Perez et al., 2022) hint at the conservation of these alternative outcomes of aldoxime metabolism in many plant species. Ultimately, these results suggest an interconnection between methionine metabolism, specialized metabolism, and plant growth which allows plants to integrate fine details of numerous metabolic pathways into their growth and developmental programs.

## Experimental Procedures

### Growth Conditions and Genetic Material

*Arabidopsis thaliana* Col-0 was used as wild-type plants. Plants were grown at 22°C ± 1°C with 16-h light/8-h dark photoperiod with fluorescent lighting intensity of 140 µE m^-2^ s^-1^. For seedings grown on Murashige and Skoog (MS) growth medium plates, seeds were sterilized with 20% (v/v) bleach containing 0.005% triton X-100 (Sigma-Aldrich, MO) for 10 min. After being washed with water four times, the seeds were cold treated at 4°C for three days before being planted on MS media containing 2% sucrose and 0.7% agar with or without supplementation of 300 µM methionine. For soil-grown plants, seeds were directly planted on soil after three days of cold treatment at 4°C. The *ref2* (*ref2-1*), *ref5* (*ref5-1*), and *cyp79b2 cyp79b3* (*b2b3*) mutants were genotyped following previously defined methods (Zhao et al., 2002; Hemm et al., 2003; Kim et al., 2015). The *b2b3ref2* triple mutant was generated by crossing *b2b3* and *ref2* plants and genotyping of the F2 progeny was done by following published methods (Zhao et al., 2002; Hemm et al., 2003).

### Plasmid Construction and Transgenic Plant Generation

To generate the Arabidopsis *CYP79F1* overexpression construct, the open reading frame of *CYP79F1* with attB sequence was synthesized from GenScript (Piscataway, NJ, USA). The synthesized DNA fragment was cloned into Gateway entry vector pCC1155 by BP cloning and subsequently recombined with the destination vector pCC0995 by LR cloning, generating the *35S:CYP79F1* construct. The *35S:CYP79F1* construct was confirmed by sequencing and was introduced into *Agrobacterium tumefaciens* (GV3101). The construct was introduced into Arabidopsis wild type or *ref2* plants via *A. tumefaciens*-mediated floral dipping method showed in Zhang et al. (2020). More than ten T1 plants were screened by application of 0.2% Basta (Rely 280, BASF, NJ).

### IAA and IAOx Purification and Quantification

IAA and IAOx purification from two-week old whole aerial parts were performed using methods described previously in Perez et al. (2021). Samples were resuspended in water and analyzed using methods and machinery described previously (Perez et al., 2021). MRM parameters of the standards (precursor m/z, fragment m/z, radio frequency (RF) lens, and collision energy) of each compound was optimized on the machine using direct infusion of the authentic standards. IAA and [^13^C_6_]-IAA were purchased from Cambridge Isotope Laboratories, and IAOx was synthesized as described previously in Perez et al. (2021). For IAA and IAOx quantification, the mass spectrometer was operated in positive ionization mode at ion spray voltage 4800V. Formic acid (0.1%) in water and 100% acetonitrile were employed as mobile phases A and B respectively with a gradient program (0-95% solvent B over 4 min) at a flow rate of 0.4 mL/min. The sheath gas, aux gas, and sweep gas were set at 50, 9, and 1 (arb unit), respectively. Ion transfer tube and vaporizer temperatures were set at 325°C and 350°C, respectively. For MRM monitoring, both Q1 and Q3 resolutions were set at 0.7 FWHM with CID gas at 1.5 mTorr. The scan cycle time was 0.8 s. MRM for IAA and IAOx was used to monitor parent ion→ product ion reactions for each analyte as follows: m/z 175.983 → 130.071 (CE, 18V) for IAA; m/z 182.091→ 136 (CE, 18V) for [^13^C_6_]-IAA; m/z 175.087→ 158 (CE, 16V) for IAOx. IAA and IAOx quantifications were conducted with four biological replicates for controls and mutants and five individual T1 plants for *F1-cos* lines.

### HPLC Analysis of Soluble Metabolites

Soluble metabolites were extracted from Arabidopsis samples using 50% methanol (v/v) incubated at 65 °C for 1 hour, with a tissue concentration of 200 mg/mL. Samples were centrifuged at 10,000 g for 10 min, and the supernatant was collected. The High-performance liquid chromatography (HPLC) analysis of metabolites was performed using an UltiMate 3000 HPLC system (ThermoFisher Scientific, MA). The system was equipped with an autosampler that was cooled to 10 °C and a diode array detector (DAD). Two columns and running methods were used to analyze different metabolites as per the specific requirements of each compound. To detect sinapoylmalate and intact indole-3-methyl glucosinolate (I3M) contents, an AcclaimTM RSLC120 C18 column (100 mm x 3 mm, 2.2 µm) (ThermoFisher Scientific, MA) was used in conjunction with mobile phases consisting of solvent A (0.1% formic acid (v/v) in water) and solvent B (100% acetonitrile) with a linear gradient of 14–18% solvent B for 10 minutes. The flow rate was set at 0.5 mL/min, and the column temperature was maintained at 40 °C. For desulfoglucosinolate quantification, samples were extracted with 50% methanol containing 250 μM sinigrin (internal standard). 100 µL of the extract was incubated with 200 µL of QAE Sephadex solution (Sigma-Aldrich, MO) for 5 minutes at room temperature. Then, the beads were washed twice with 50% methanol and twice with autoclaved MilliQ water. After the final wash, 100 µL of MilliQ water containing sulfatase (Sigma-Aldrich, MO) were added to the samples, which were then incubated at 37 °C for 6 hours. 10 µL of desulfied samples were analyzed using the HPLC equipped with an AcclaimTM 120 C18 column (150 mm x 4.6 mm, 5 µm) (ThermoFisher Scientific, MA). Metabolites were separated by utilizing a mobile phase composed of solvent A (water) and solvent B (100% acetonitrile), and a linear gradient program of solvent B 2-12% over 10 minutes, 12-15% over 15 minutes, 15-25% over 17.5 minutes, and 95% for 2 minutes. The flow rate was set at 1 mL/min, and the column temperature was maintained at 40 °C. The content of sinapoylmalate was quantified by measuring the peak area at 328 nm and comparing it to that of sinapic acid (Sigma-Aldrich, MO) equivalents. The content of desulfo-glucosinolates was quantified using the peak area at 220 nm and response factors (Brown et al., 2003).

### Gene Expression Analysis

Total RNA was extracted from young rosette leaves and inflorescence stem of four-week-old plants using the TRIzol method as per the manufacturer’s protocol (ThermoFisher Scientific, MA). Two μg of total RNA were subjected to reverse transcription using the High-Capacity cDNA Reverse Transcription Kit (ThermoFisher Scientific, MA) with oligo(dT) at 65°C for 2 hr. PCR was performed as described previously (Hansen et al., 2001, Chen et al., 2003). The following specific conditions were used: *CYP79F1* experiment performed with 53 °C and 35 cycles, *CYP79F2* experiment performed with 53 °C and 40 cycles, and *TUB3* experiment performed with 55 °C and 35 cycles. The sequences of the primers are included in Table S1. The PCR products were analyzed by agarose gel electrophoresis.

### PAL Activity Test

The PAL activity was measured using the protocol outlined in Kim et al., (2015) with certain modifications. Crude proteins were extracted from the frozen 5^th^ to 7^th^ true leaves of four-week-old Arabidopsis plants by grinding them completely using a Benchmark BeadBlaster 24 homogenizer (Benchmark Scientific, NJ) and then mixing the powder with an extraction buffer (0.1 M Tris-HCl (pH 8.3), 10% glycerol, and 5 mM DTT). Crude protein concentration was determined using the Bradford Reagent (Sigma-Aldrich, MO), following manufacturer’s instructions. The enzyme reaction of PAL was started by adding 150 µL of crude protein to 400 µL of reaction buffer containing 5 mM phenylalanine. The reaction mixture was then incubated at 37 °C for 90 minutes. The reaction was terminated by adding 40 µL of 30% (v/v) acetic acid. The reaction products were extracted with 600 µL of ethyl acetate, followed by evaporation of the extracts using an Eppendorf Vacufuge Plus (Eppendorf, Hamburg, Germany). The dried extract was then redissolved in 100 µL of 50% methanol and 10 µL of extract was analyzed using the HPLC with a linear gradient of solvent B 12–30% for 2.6 minutes, 30–95% for 4 minutes, and 95% for 3 minutes. The flow rate was set at 0.7 mL/min, and the column temperature was maintained at 40 °C. The content of *trans*-cinnamic acid, the reaction product, was quantified by measuring the peak area at 270 nm and comparing it to that of authentic standard *trans*-cinnamic acid (Sigma-Aldrich, MO). PAL activity assays were performed with four biological replicates for *ref2, F1-OX/ref2*, and *F1-cos/ref2*, and two biological replicates for wild type. The statistical significance of the results was tested by one-way ANOVA.

### Methionine extraction and detection

Whole aerial parts of 3-week-old plants were used for methionine quantification using the extraction method slightly modified from the previously described method (Cao et al., 2019). The plant materials were extracted with the extraction buffer containing 2 μM labeled Met (13C, 15N, Cambridge Isotope Laboratories), 10 μM DL-dithiothreitol (DTT, Gold Biotechnology), and 10 mM perfluoroheptanoic acid (PFHA, Sigma-Aldrich) at 90°C for 10 min at a tissue concentration of 100 mg/mL. Then, the extracts were centrifuged at 13,000 g at 4°C for 10 min. 100 µL of extracts were filtered on polytetrafluoroethylene (PTFE) filter (Millipore) and the final elutes were fully dried in the vacuo and stored at −20°C until further analysis. All samples were resuspended in 500 µL of water.

The compounds were analyzed using the triple quadrupole TSQ Altis mass spectrometers (Thermo Fisher Scientific, San Jose, CA, USA) coupled with the Vanquish Horizon UHPLC (Thermo Fisher Scientific). The TSQ Altis was housed with a heated-electrospray ionization (HESI) source using the following source settings with sheath gas flow with 50 Arb, auxiliary gas flow with 10 Arb, sweep gas flow with 1 Arb, ion transfer tube temperature at 325 °C, and vaporizer temperature at 350 °C. The spray voltage was 4.0 kV under the positive polarity. A scan time was set at 1 sec and the Q1 and Q3 resolutions of full width at half maximum (FWHM) were both 0.7. For the CID gas pressure, 1.5 mTorr was used. To determine the optimal fragments and collision energies (CE) for MRM transitions, we utilized the Xcalibur 4.1 software from Thermo Fisher Scientific for optimization. The selected fragment ions of the methionine precursor ion (150 m/z) were observed at 56, 104, and 132.917 m/z, with collision energies set at 19, 12, and 11 V, respectively. The corresponding fragment ions of the isotopically labeled form for L-methionine-13C5,15N (156 m/z) were detected at 60, 109, and 138 m/z with the collision energies set at 20, 13, and 11 V, respectively. The optimal quantification optimization method of methionine was best achieved using the fragment ion at 132.917 m/z, while for 13C5 methionine, the optimal fragment ion was also observed at 138 m/z. The compounds were separated using Atlantis T3 3 µm, 2.1 mm X 100 mm (P/N 186003718; Waters, Milford, MA, USA). To accomplish LC separation, a Vanquish UHPLC system was set for the column compartment temperature at 40 °C, and the flow rate was set to 125 μL/min. The mobile phases consisted of the mobile phase A (99.9% water (v/v), and 0.1% formic acid (v/v)) and the mobile phase B (99.9% acetonitrile (v/v), and 0.1% formic acid (v/v)). The following linear gradient was applied from 0% to 10% solvent B in 3.6 min, ramping up to 80% in 4 min, holding 80 % solvent B in 1 min, and ramping down 0% solvent B in 0.2 min, and staying 0% solvent B for 7 min as a re-equilibration. The samples were kept at 6 °C in the autosampler. The injection volume was 1 μL. Xcalibur 4.1 software (Thermo Fisher Scientific) was used to determine the compounds and processed the quantification using the Quant Brower. To obtain the absolute expression value of the metabolite, the peak area of the compound is normalized by the initial amount of sample and the input amount of the isotopic methionine. This normalized peak area is then compared to the standard curve of the metabolite standards, allowing for the determination of its quantitative value.

### Accession Numbers

*CYP79F1* (At1g16410)

*CYP79F2* (At1g16400)

*CYP79B2* (At4g39950)

*CYP79B3* (At2g22330)

*CYP83A1/REF2* (At4g13770)

*CYP83B1/REF5* (At4g31500)

## Author Contributions

D.S., V.C.P, and J.K. designed the research project; G.K.D., D.S., V.C.P, R.D., H.Z., N.Z., J.K., and K.H.C. performed the experiments; B.T. and A.G. synthesized aldoximes; and analyzed the data; D.S., V.C.P., and J.K. wrote the manuscript. All authors read and agreed with the manuscript.

## Acknowledgments

This work was supported by National Science Foundation CAREER Grant (IOS-2142898), and a startup fund from the Horticultural Sciences Department and Institute of Food and Agricultural Sciences at the University of Florida to J.K. National Institutes of Health (NIH)-5R35GM137893-03 to AG.

## The List of Supporting Information

**Figure S1.** Expression of *CYP79F1* and *CYP79F2* in wild type, *ref2, F1-cos* plants and non-bushy *CYP79F1* overexpression plants in the wild-type and *ref2* genetic backgrounds.

**Figure S2.** Polyamine and ACC treatment did not phenocopy morphology of *F1-cos* lines.

**Table S1.** List of primers

