## Supplemental Data for "Altered methionine metabolism impacts phenylpropanoid production and plant development in *Arabidopsis thaliana*"

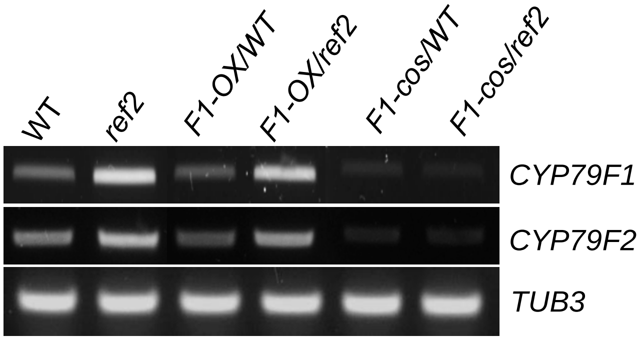


**Figure S1.** Expression of *CYP79F1* and *CYP79F2* in wild type, *ref2, F1-cos* plants and non-bushy *CYP79F1* overexpression plants in the wild-type and *ref2* genetic backgrounds.

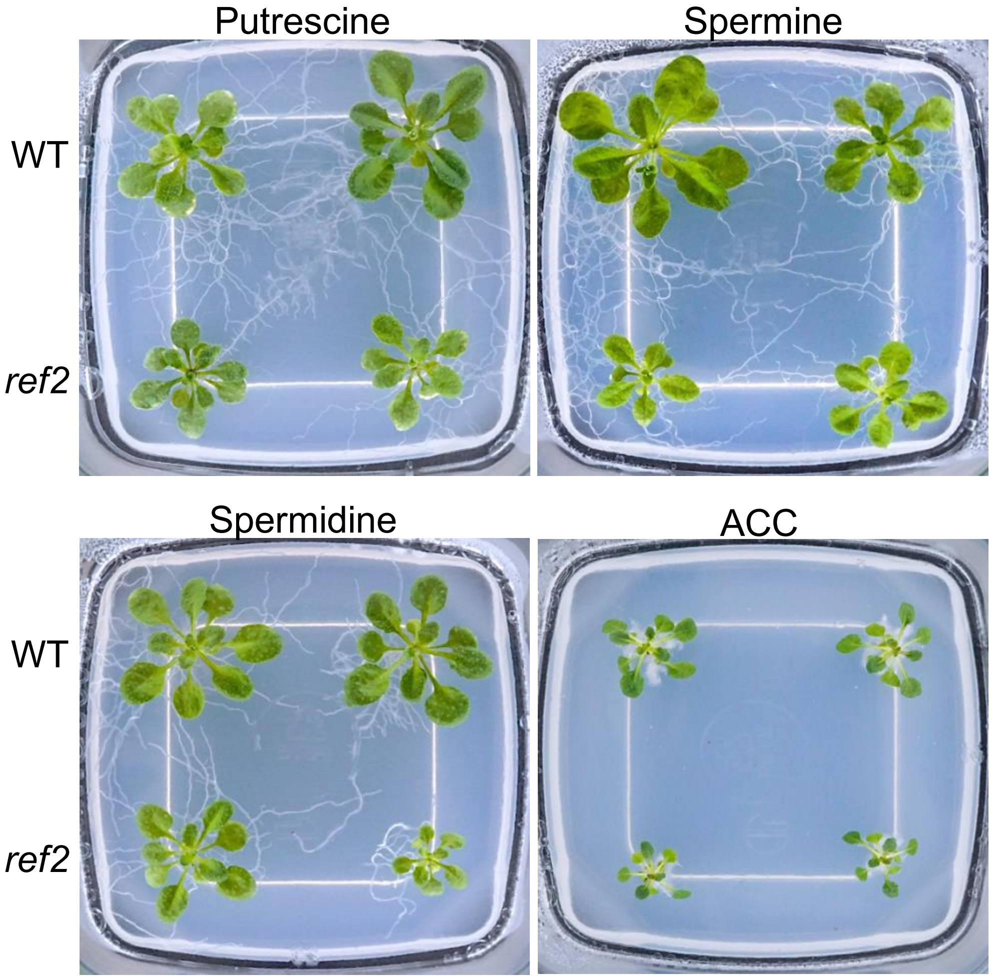


**Figure S2. Polyamine and ACC treatment did not phenocopy growth morphology of *F1-cos* lines.** Representative images of wild type and *ref2* grown on polyamines (putrescine, spermine, and spermidine) and the ethylene precursor 1-aminocyclopropane-1-carboxylate (ACC) for three weeks. Plants grown with supplement of 500 μM of polyamines or 30 μM of ACC on growth media did not show any growth changes similar to those seen in *F1-cos* lines.

**Table S1.** List of primers

| Primer Name | Sequence (5’->3’) |
|---|---|
| TUB3-F | TGGTGGAGCCTTACAACGCTACTT |
| TUB3-R | TTCACAGCAAGCTTACGGAGGTCA |
| CYP79F1 RT-Forward | AAAGCTCAATGCGTAGAAT |
| CYP79F1 RT-Reverse | TTTTTAGACACCATCTTGTTTTCTTCTTC |
| CYP79F2 RT-Forward | AAAGCTCAATGCGTCGAAT |
| CYP79F2 RT-Reverse | GCGTCGAAACACATCACAGAG |
